# Rapid degradation of GRASP55 and GRASP65 reveals their immediate impact on the Golgi structure

**DOI:** 10.1101/2020.07.07.192609

**Authors:** Yijun Zhang, Joachim Seemann

## Abstract

GRASP65 and GRASP55 have been implicated in stacking of Golgi cisternae and lateral linking of stacks within the Golgi ribbon. However, loss of gene function approaches by RNAi or gene knockout to dissect their respective roles often resulted in conflicting conclusions. Here, we gene-edited GRASP55 and/or GRASP65 with a degron tag in human fibroblasts, allowing for the induced rapid degradation by the proteasome. We show that acute depletion of either GRASP55 or GRASP65 does not affect the Golgi ribbon, while chronic degradation of GRASP55 disrupts lateral connectivity of the Golgi ribbon. Acute double depletion of both GRASPs coincides with the loss of the vesicle tethering proteins GM130, p115 and Golgin-45 from the Golgi and compromises ribbon linking. Furthermore, neither GRASP55 and/or GRASP65 are required for maintaining stacks or *de novo* assembly of stacked cisternae at the end of mitosis. These results demonstrate that both GRASPs are dispensable for Golgi stacking, but are involved in maintaining the integrity of Golgi ribbon together with GM130 and Golgin-45.

## Introduction

The Golgi apparatus is a membrane-bound organelle essential for processing and sorting of secretory and membrane proteins. As the central module of the exocytic pathway, it receives proteins and lipids from the endoplasmic reticulum, post-translationally modifies and sorts them to their appropriate cellular destinations (Pantazopoulou and Glick, 2019). This secretory function of the Golgi is highly conserved among eukaryotes and is carried out by stacks of flattened, disk-shaped cisternae. Concentrating cisternae into spatially proximate stacks enables efficient cargo sorting between adjacent cisternae as well as post-translational modifications such as glycosylation (Lowe, 2011). Most vertebrate cells contain 100-200 stacks, each comprised of four to six cisternae (Storrie et al., 2012; Sin and Harrison, 2016). In addition, these stacked cisternae are laterally interconnected into a single, contiguous ribbon-like entity that is positioned next to the centrosomes in the perinuclear region (Lowe, 2019). By joining adjacent cisternae between stacks, the Golgi can accommodate and process large secretory cargos that do not fit into individual cisternae such as collagen or Weibel-Palade bodies (Ferraro et al., 2014; Lavieu et al., 2014). On the other hand, the concentration and the polarized positioning of the Golgi ribbon within the cells facilitates the directional delivery of cargo and signaling molecules and is thereby indispensable for cell polarization and differentiation (Wei and Seemann, 2017).

Despite the characteristic organization and unique morphological appearance, Golgi membranes are highly dynamic and are constantly remodeled to accommodate extensive vesicular trafficking through the stacks (Lowe, 2011). An imbalanced flux of membranes through the Golgi leads to fragmentation of the Golgi ribbon, which is commonly observed under pathological conditions (Wei and Seemann, 2017; Makhoul et al., 2019). Physiologically, this process is harnessed during mitosis, where a block of vesicle fusion initiates the conversion of the Golgi ribbon into vesicles. Disassembly of the ribbon is not only required for mitotic progression (Sütterlin et al., 2002; Colanzi et al., 2007; Guizzunti and Seemann, 2016), it also facilitates Golgi partitioning by the spindle into the daughter cells, where vesicles fuse to reform a ribbon of stacked cisternae (Shorter and Warren, 2002; Wei and Seemann, 2009a). Such dramatic structural rearrangements raise a tremendous interest to investigate the proteins and the molecular mechanisms underlying Golgi stacking and lateral linking.

The highly dynamic and unique structure of the Golgi critically depends on structural Golgi proteins, which jointly act as a scaffold or matrix to support its morphology and function (Barinaga-Rementeria Ramirez and Lowe, 2009; Gillingham and Munro, 2016). These structural Golgi proteins are peripherally associated with the cytoplasmic face of the Golgi membrane and are categorized into two major protein families: GRASPs and Golgins. GRASPs (Golgi reassembly and stacking proteins), including GRASP55 and GRASP65, were first identified by their roles in post-mitotic stacking of cisternae in a cell-free system (Barr et al., 1997; Shorter et al., 1999). On the other hand, Golgins, such as GM130, Golgin-97, Golgin-84 and Golgin-45, are long rod-like coiled-coil proteins that act as vesicle tethers at distinct cisternae to facilitate cargo transport through the stack (Gillingham and Munro, 2016). Notably, members of the GRASP and Golgin families form distinct subcomplexes with each other. For example, GRASP65 is recruited to the *cis*-Golgi by GM130 while GRASP55 and Golgin-45 form a complex at *medial*/*trans*-Golgi cisternae (Barr et al., 1998; Short et al., 2001). Furthermore, GRASP55 and GRASP65 also directly participate in vesicle transport through the Golgi by binding to p24 cargo receptors (Barr et al., 2001).

The role of GRASP55 and GRASP65 in establishing and maintaining the integrity of the Golgi ribbon is thought to depend on their ability to physically bridge apposing cisternae. Both GRASPs share a highly homologous GRASP domain at their N-termini, which allows each GRASP to assemble into anti-parallel homo-oligomers in *trans* (Wang et al., 2005; Feng et al., 2013). The oligomerization of GRASP65 is sufficient to hold adjacent membrane compartments together, as demonstrated by clustering of mitochondria induced by expression of GRASP65 fused to a mitochondria targeting sequence (Bachert and Linstedt, 2010). Based on their membrane linking capacity, GRASP55 and 65 have been implicated in packing cisternae into stacks as well as in lateral linking adjacent stacks into a ribbon. During mitosis, phosphorylation of GRASP55 and 65 breaks the *trans*-oligomers, which coincides with lateral unlinking of the ribbon and unstacking of the cisternae (Wang et al., 2003; Tang et al., 2010). However, the precise function of GRASPs in organizing the Golgi structure *in vivo* is still unclear.

In efforts to dissect the contributions of GRASPs to Golgi stacking and ribbon linking, several RNAi and gene knockout-based studies have been conducted in both cellular and animal models, which led to inconsistent results and conclusions. Downregulation of either GRASP55 or 65 by RNAi were reported to only induce Golgi fragmentation or ribbon unlinking (Sütterlin et al., 2005; Duran et al., 2008; Feinstein and Linstedt, 2008; Puthenveedu et al., 2006), while others reported a loss of cisternae (Xiang and Wang, 2010; Bekier et al., 2017). Similarly, double suppression of GRASP55 and 65 by RNAi or CRISPR-mediated gene inactivation disrupted the stacks and vesiculated the cisternae (Xiang and Wang, 2010; Bekier et al., 2017), while others reported no effect on stacking upon RNAi and in knockout mice (Lee et al., 2014; Grond et al., 2020).

One key limitation of these RNAi or gene knockout-based approaches is that protein levels are gradually reduced over an extended period of time. Typically, it takes 2-3 days or sometimes up to weeks to efficiently eliminate protein expression. As such, chronic depletion might lead to complex phenotypes that are indirect and difficult to interpret. In addition, long-term loss of critical functions provides cells ample time to adapt and skew the phenotypes by activating compensatory pathways. To avoid such chronic complications, we set out to determine the immediate effects caused by acute loss of GRASP55 and 65 at the protein level. To this end, we employed the auxin-inducible degron (AID) system (Nishimura et al., 2009; Holland et al., 2012) to acutely degrade GRASP55 and/or GRASP65 within two hours. Our results show that simultaneous depletion of both GRASPs, but not individually, displaces GM130 and p115 from the Golgi and indirectly affects the lateral linking of the Golgi ribbon. Notably, consistent with the ribbon unlinking effect reported for RNAi, long-term degradation of GRASP55 is also sufficient to disrupt the Golgi ribbon, while short-term degradation does not. Moreover, acute elimination of GRASP55 and/or GRASP65 did not affect Golgi stacking during interphase or in cells that reassembled a Golgi ribbon at the end of mitosis.

## Results

### Generation of SV589 cell lines endogenously expressing degron-tagged GRASP55 and/or GRASP65

To study the cellular effects caused by acute loss of GRASP55 and GRASP65 (briefly referred to as GRASP55 and 65), we employed the auxin-inducible degron (AID) technology. When cells are incubated with the small molecule auxin, proteins with an AID tag are rapidly degraded by the proteasome (Nishimura et al., 2009; Holland et al., 2012). Degradation of AID-tagged proteins requires expression of the auxin perceptive F-box protein TIR1 from *Oryza sativa* (OsTIR1), which forms a functional E3 ubiquitin ligase complex with endogenous SCF (Skp1–Cullin–F-box) of the host cells. When added to cells, the auxin Indole-3-acetic acid (IAA) binds to TIR1, leading to polyubiquitination and acute proteolysis of target proteins tagged with the minimal AID (mAID) sequence of 68 amino acids (Kubota et al., 2013). To create homozygous cell lines where endogenous GRASP55 and/or 65 are tagged with mAID (hereafter referred to as RC55, RC65 and RC65+55 cell lines, respectively), we employed the CRISPaint (CRISPR-assisted insertion tagging) system (Schmid-Burgk et al., 2016). Using this approach, a double-strand break was induced in front of the stop codon where a repair sequence was then integrated. This exchanges the endogenous stop codon with the 3xFlag epitope tag (to facilitate detection), followed by the mAID degron, a T2A self-cleaving peptide and an antibiotic resistance gene (Fig. 1A and B). We successfully established immortalized SV589 human fibroblast cell lines that endogenously express either GRASP55-3xFlag-mAID (RC55), GRASP65-3xFlag-mAID (RC65) or both (RC65+55, generated by endogenous tagging GRASP55 with 3xFlag-mAID in RC65 cells).

**Figure 1.**
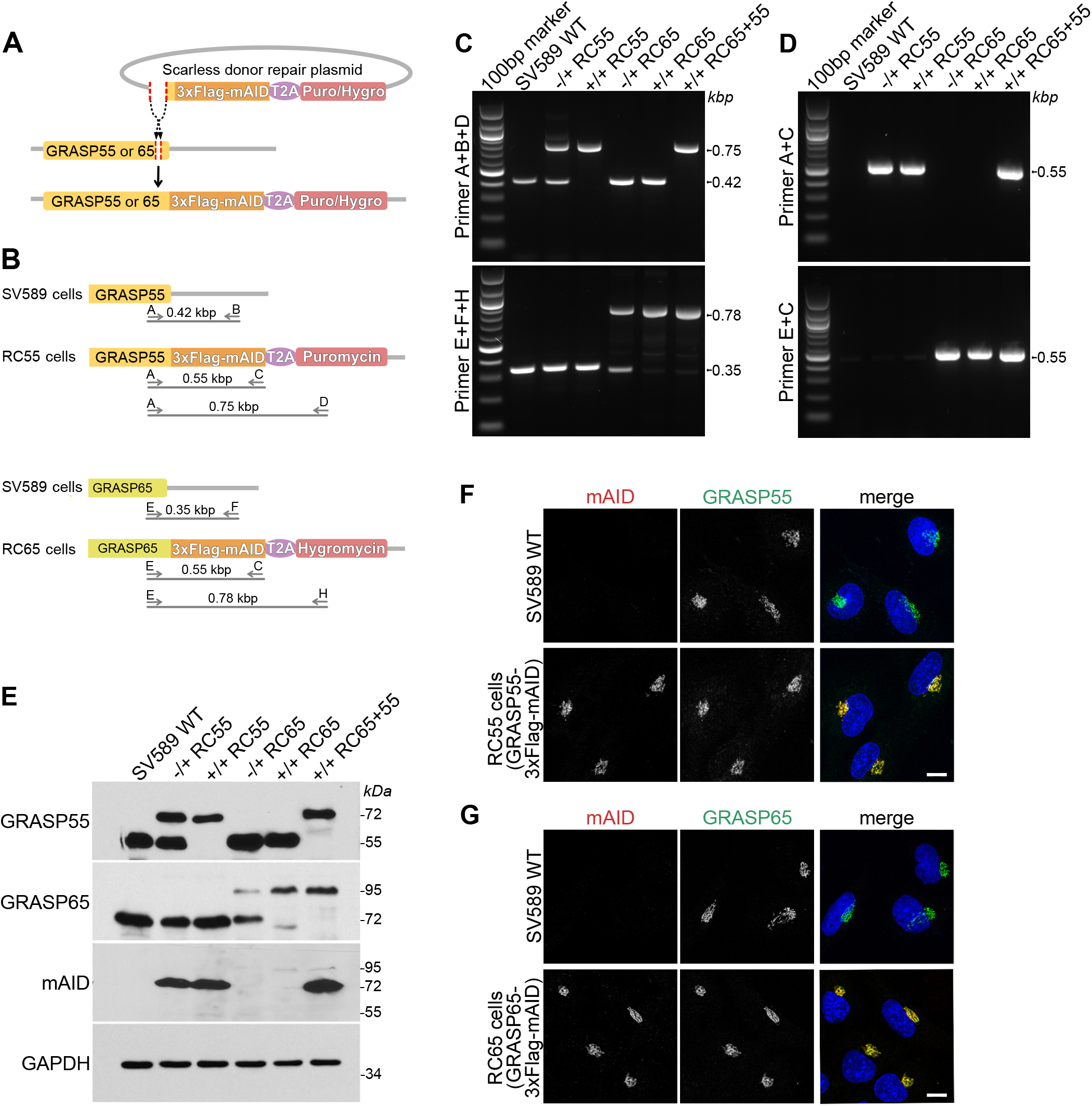
Generation of cell lines expressing endogenous GRASP55 and/or GRASP65 tagged with the minimal auxin inducible degron (mAID) A). Scheme for tagging the endogenous loci of GRASP55 or GRASP65 with 3xFlag and mAID. The DNA sequence encoding 3xFlag-mAID separated by a self-cleaving T2A peptide from the resistance gene (puromycin or hygromycin) were inserted by CRISPR-Cas9 gene editing in front of the stop codon of GRASP55 or GRASP65. The amino acids between the insert location and the stop codon were scarlessly repaired by adding the sequence in front of 3xFlag-mAID of the repair donor plasmid. B). Scheme of the primer sets used for genotyping gene edited cell lines. The cell line with GRASP55 scarlessly tagged with 3xFlag-mAID and puromycin is referred to as RC55, GRASP65 scarlessly tagged with 3xFlag-mAID and hygromycin as RC65, and both GRASP55 and 65 tagged with 3xFlag-mAID and indicated resistance genes as RC65+55. C, D). Genomic PCR to genotype heterozygous (-/+) and homozygous (+/+) RC55, RC65 and PC65+55 cell lines for GRASP55 tagged with 3xFlag-mAID or GRASP65-3xFlag-mAID. Parental SV589 cells are shown as the negative control (WT). E). Immunoblot analysis of whole cell lysates of heterozygous (-/+) and homozygous (+/+) RC55, RC65 and RC55+65 cells using antibodies against GRASP55, GRASP65, the degron tag mAID and GAPDH. F). Parental SV589 cells and RC55 cells expressing GRASP55-3xFlag-mAID were immunostained for mAID (red), GRASP55 (green) and labeled for DNA (blue). G). SV589 cells and RC65 cells stably expressing GRASP65-3xFlag-mAID were immunostained for mAID (red), GRASP65 (green) and labeled for DNA (blue).

We identified heterozygous and homozygous clones by genomic PCR using forward primers targeting the 3’-region of the GRASP55 or 65 coding sequences (primers A and E) and reverse primers binding to the 3’-UTRs of each GRASP (primers B and F), the mAID sequence (primer C), or the antibiotic resistance gene (primers D and H), respectively (Fig. 1B-D). We then performed western blotting analysis to confirm that the cell lines indeed expressed mAID-tagged GRASP proteins. Compared to parental SV589 cells, the bands recognized by antibodies against GRASP55 or 65 were upshifted (corresponding to mAID and Flag tag) in RC55 and RC65 cells and were also detected by an anti-mAID antibody (Fig. 1E). Of note, the mAID band corresponding to GRASP65-3xFlag-mAID at 95 kDa was much weaker compared to the GRASP55-3xFlag-mAID signal at 72 kDa, indicating that the endogenous protein levels of GRASP55 are much higher than GRASP65. This result is consistent with a previous quantitative proteomics report showing that the protein copy number of GRASP55 in HeLa cells is 45 fold higher than that of GRASP65 (Kulak et al., 2014). In accordance with the western blotting results, our immunofluorescence analysis revealed that mAID can only be detected in RC55, RC65 or RC55+65 cells, but not in parental SV589 cells (Fig. 1F, G and Fig. S1C). Furthermore, the staining for mAID and GRASP55 or 65 localized to the Golgi ribbon in the perinuclear region, demonstrating that the C-terminal 3xFlag-mAID-tag does not alter Golgi targeting of the GRASP proteins (Fig. 1F and G).

### GRASP55 and GRASP65 are acutely degraded within two hours upon auxin induction

To enable acute degradation of mAID-tagged proteins, we stably introduced myc-tagged and codon-optimized OsTIR1 (TIR1-2xMyc) into homozygous RC55, RC65 and RC65+55 cells. In line with previous studies (Natsume et al., 2016), we noticed that constitutive expression of TIR1 reduced the protein levels of mAID-tagged proteins even in the absence of IAA (data not shown). To circumvent potential side effects caused by chronic basal degradation, we generated RC55, RC65 and RC65+55 cell lines expressing TIR1-2xMyc under doxycycline control to precisely induce its timely expression. Since only homozygous RC55, RC65 and RC65+55 cells stably expressing inducible TIR1-2xMyc were used for further experiments, for simplicity we hereafter refer to them as RC55, RC65 and RC65+55 cells. We first compared the stability of mAID-tagged GRASP55 and 65 after short-term 6 h induction or long-term 20 h induction of TIR1. 20 h TIR1 expression in the absence of IAA significantly decreased GRASP55 and 65 levels (Fig. S1A). By contrast, 6 h TIR1 induction showed only minimal to no basal degradation of GRASP55 and 65 (Fig. 2A). A subsequent treatment with IAA led to complete degradation of both GRASPs within 2 hours but did not change the protein levels of the Golgi resident proteins Golgin-84 or the recruitment of the GM130-associated tethering factor p115 (Fig. 2A and S1A).

**Figure 2.**
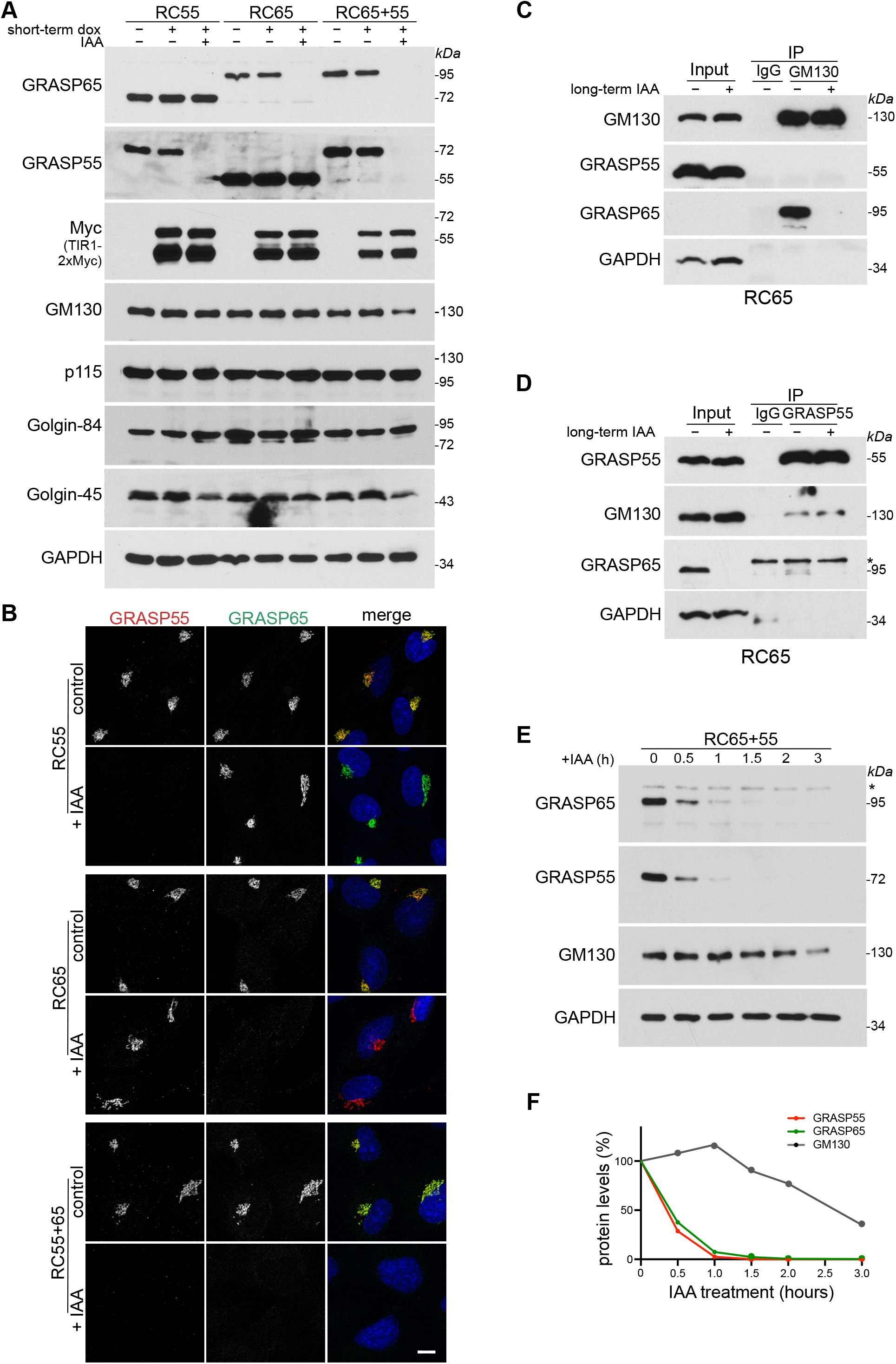
The auxin IAA induces rapid depletion of mAID-tagged GRASP55 and GRASP65. A). Homozygous RC55, RC65 and RC65+55 cells inducibly expressing TIR1-2xMyc were treated with doxycycline for 6 h (short-term dox) and then with the auxin indole-3-acetic acid (IAA) for a further 2 h to degrade GRASP55 and/or 65. Cell lysates were immunoblotted with indicated antibodies. B). RC55, RC65 and RC65+55 cells were doxycycline treated for 6 h and with IAA for a further 2 h as in (A), immunolabelled for GRASP55 (red), GRASP65 (green) and stained for DNA (blue). Scale bar, 10 μm. C, D). GRASP55 does not compensate for the loss of GRASP65 to stabilize GM130. RC65 cells were treated with doxycycline for 6 h before adding IAA for 2 days to deplete GRASP65. GM130 (panel C) and GRASP55 (panel D) were then immunoprecipitated from the cell lysates and analyzed by western blotting. * denotes unspecific band. E). Simultaneous degradation of GRASP55 and 65 causes a delayed reduction of GM130 levels. RC65+55 cells expressing TIR1 for 6 hours were treated with IAA. Cells lysates were collected at the indicated time points and subjected to western blotting with the indicated antibodies. * denotes unspecific band. F). Graph showing relative protein levels in (E) at different time points.

Degradation of GRASP55 and 65 was further corroborated by means of immunofluorescence microscopy. Short-term GRASP55 elimination did not alter the signal of GRASP65 on the Golgi and vice versa (Fig. 2B), and depletion of both GRASPs had no effect on the localization of the GFP-tagged Golgi enzyme *N*-acetylglucosaminyl transferase I (NAGTI-GFP) (Fig. S1B). Furthermore, after 2 h IAA treatment the mAID signal was lost from the Golgi, while the perinuclear localization of Golgin-84 remained (Fig. S1C). Together, these results indicate that acute depletion of GRASP55 and 65 does not disperse the Golgi ribbon, suggesting that neither GRASP55 nor 65 is required for maintaining the perinuclear Golgi morphology.

Next, we determined whether degradation of GRASP55 and 65 affects the stability of their respective binding partners. Both GRASP proteins form tight complexes via their PDZ domains with members of the Golgin protein family. Specifically, GRASP55 binds to Golgin-45, a *medial*-Golgin that facilitates secretory protein transport (Short et al., 2001), while GRASP65 interacts with GM130, a *cis*-Golgi matrix protein that facilitates vesicle docking by forming a complex with p115 (Barr et al., 1998). Whether degradation of GRASP55 and 65 affects the stability of Golgin-45 and GM130 was tested by western blotting. Surprisingly, while Golgin-45 was notably reduced upon degradation of GRASP55 (Fig. 2A), GM130 was not affected when either GRASP55 or 65 was depleted, showing that GRASP65 is not degraded in a complex with GM130. However, GM130 levels decreased when both GRASPs were simultaneously eliminated (Fig. 2A and E), which was also observed after chronic down-regulation by siRNA (Xiang and Wang, 2010; Lee et al., 2014). One possible explanation might be that once GRASP65 is degraded, GRASP55 compensates for the loss of GRASP65 and binds via its GRASP domain to stabilize GM130, which was reported for *in vitro* translated GRASP55 and GM130 (Shorter et al., 1999). To test if GRASP55 interacts with GM130 in cells, we degraded GRASP65 in RC65 cells for two days and then immunoprecipitated GM130 or GRASP55. GRASP65 co-pelleted with GM130 from control lysates as reported (Barr et al., 1997), but GRASP55 was neither pulled down by GM130 from control cells nor from GRASP65 depleted lysates (Fig. 2C). As previously reported (Short et al., 2001), we also found a minor fraction of GM130 co-immunoprecipitated with GRASP55, but GRASP55 was below the detection limit in GM130 immunoprecipitates (Fig. 2C and D). However, no increase in the amount of GM130 pulled down by GRASP55 was detected after GRASP65 was depleted, demonstrating that GRASP55 does not compensate for the loss of GRASP65 to stabilize GM130.

To further evaluate GM130 stability upon acute depletion of GRASP55 and 65, we followed the degradation over a time course (Fig. 2E and F). The protein levels of GRASP55 and 65 in IAA-treated RC65+55 cells were significantly decreased within 30 min, and only trace amounts remained after 1 hour. On the other hand, GM130 levels appeared to be unaffected after 1 h, but started to decrease by 2 h and were significantly diminished after 3 hours. The delay of GM130 loss compared to GRASP55 and 65 suggests that the stability of GM130 was indirectly compromised by the simultaneous loss of GRASP55 and 65. Considered together, the data shown in Fig. 2 suggest that induced degradation is highly efficient in our system and that the degradation of GRASP55 and/or 65 affect the stability of Golgin-45 and GM130 but does not change the overall morphology of the perinuclear Golgi ribbon.

### Acute double degradation of GRASP55 and GRASP65 partially displaces GM130 and p115 from the Golgi

Because depletion of GRASP55 and 65 coincided with a time-delayed reduction of GM130 levels, we analyzed whether it also alters the localization of GM130. Cells were treated with IAA for 2 h to degrade GRASP55 and/or 65 and then double-stained for GM130 and another *cis*-Golgi marker Golgin-84. GM130 and Golgin-84 co-localized at perinuclear Golgi ribbon in control cells, which was not changed after the degradation of either GRASP55 or 65 (Fig. 3A). By contrast, in cells depleted of both GRASPs, the overlapped intensity of GM130 with Golgin-84 was reduced and GM130 was partially displaced from the perinuclear Golgi ribbon. In line with our previous report, Golgi-displaced GM130 re-localized to the nucleus via its N-terminal NLS, which became more noticeable after increasing the brightness of the image (Fig. 3A) (Wei et al., 2015). In addition, since p115 is recruited to the Golgi by GM130 (Nakamura et al., 1997; Seemann et al., 2000a), we determined whether p115 Golgi localization was affected by depletion of GRASPs (Fig. 3B). Co-localization of p115 and GM130 at the Golgi was unaffected when either GRASP55 or 65 was degraded. In marked contrast, in IAA-treated RC55+65 cells, p115 was displaced together with GM130 from the perinuclear region where the Golgi-resident enzyme NAGTI-GFP remained. Unlike GM130, p115 was redistributed from the Golgi to the cytosol and not detected in the nucleus (Fig. 3B), and its protein level was not reduced either (Fig. 2A, Fig. 3C).

**Figure 3.**
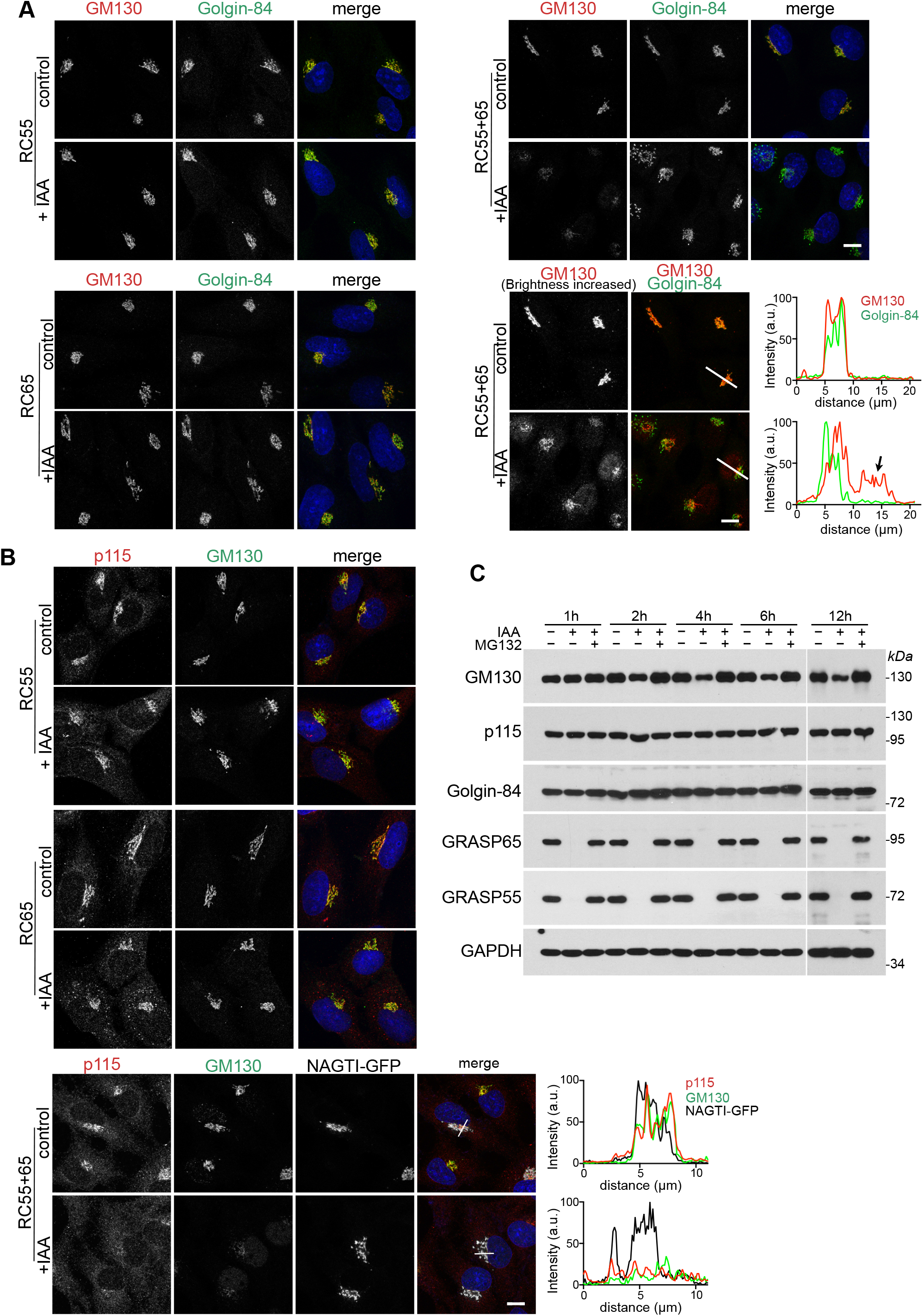
Acute simultaneous depletion of GRASP55 and 65, but not separately, partially dislocates GM130 from the Golgi. A). GM130 is partially displaced from the Golgi and relocated to the nucleus after degradation of GRASP55 together with 65, but not after individual depletion of GRASP55 or 65. RC55, RC65 or RC65+55 cells were treated with doxycycline for 6 h and then with IAA for a further 2 h. The cells were then fixed and immunolabelled for GM130 (red), the cis-Golgi protein Golgin-84 (green) and stained for DNA (blue). The brightness of the GM130 image of RC65+55 cells was increased to better visualize the redistribution of GM130. The white lines show the position of the line-scan that was used to measure the fluorescence intensity of GM130 (red) and Golgin-84 (green) across the Golgi and nucleus as shown in the graphs. Scale bar, 10 μm. B). Degradation of GRASP55 and 65 partially delocalizes p115 together with GM130 from the Golgi. RC55, RC65 and NAGTI-GFP-expressing RC65+55 cells were treated as in (A). Fixed cells were immunostained for p115 (red), GM130 (green) and labelled for DNA (blue). The Golgi is marked by the GFP fluorescence signal of the Golgi enzyme NAGTI-GFP. The white lines show the position of the line-scan that was used to measure the fluorescence intensity of p115 (red), GM130 (green) and NAGTI-GFP across the Golgi as shown in the graphs. Scale bar, 10 μm. C). GM130 is partially degraded by the proteasome after acute double depletion of GRASP55 and 65. RC65+55 cells were treated with doxycycline for 6 h, followed by IAA or IAA plus 10 μM MG132. Cell lysates were collected at the indicated time points and subjected to immunoblotting with the indicated antibodies.

Moreover, western blotting analysis showed that within 1 h of IAA addition, GRASP55 and 65 amounts dropped below detection levels, while GM130 more gradually decreased within the first 4 h, and then persisted at a stable level for the duration of the experiment of 12 hours (Fig. 3C). Blocking GRASP55 and 65 degradation with the proteasome inhibitor MG132 also stabilized GM130 and led to its accumulation. By contrast, p115 and Golgin-84 levels were not notably changed after long-term depletion of both GRASPs. Taken together, these results show that acute and simultaneous depletion of GRASP55 and 65, but not individually, partially delocalizes GM130 from the Golgi leading to its degradation by the proteasome.

### Acute degradation of GRASP55 and GRASP65 disrupts the lateral linking of the Golgi ribbon

Previous studies using RNAi and gene knockout have implicated GRASP55 and 65 in laterally connecting adjacent stacks to facilitate proper distribution of Golgi-resident enzymes within the interconnected Golgi ribbon (Puthenveedu et al., 2006; Feinstein and Linstedt, 2008; Xiang and Wang, 2010; Veenendaal et al., 2014). To determine if acute depletion of GRASP55 and/or 65 affects the lateral linking of the ribbon, we stably expressed the Golgi enzyme NAGTI-GFP in RC55, RC65 and RC65+55 cells and then performed fluorescence recovery after photobleaching (FRAP) to analyze GFP recovery as a measure of the lateral mobility within the Golgi ribbon. In contrast to previous reports using RNAi (Puthenveedu et al., 2006; Feinstein and Linstedt, 2008; Xiang and Wang, 2010), acute degradation of either GRASP55 or 65 did not change the rate of recovery, showing that the stacks remained laterally linked (Fig. 4A-C). One possibility is that GRASP55 and 65 might be required for the initial connection of the stacks during Golgi ribbon formation but are dispensable once the ribbon is established. To test this idea, cells depleted of GRASPs were first treated with nocodazole to disperse the Golgi ribbon into isolated stacks. Nocodazole was then washed out to allow the reformation of the perinuclear Golgi ribbon (Fig. 4A and S2). The rate of NAGTI-GFP recovery in cells lacking GRASP55 or 65 was indistinguishable to control cells, indicating that an interconnected Golgi ribbon reformed (Fig. 4B and C). However, acute and simultaneous degradation of both GRASPs disrupted the lateral linking of the Golgi ribbon, both in a pre-existing ribbon and in a newly reformed ribbon after nocodazole washout (Fig. 4B and C). In contrast to acute loss of GRASP55, we also noticed that chronically diminished GRASP55 levels disrupted the integrity of the ribbon, which is consistent with the previous RNAi experiments (Xiang and Wang, 2010) (Fig. 4D). Moreover, although RNAi of GRASP65 was reported to compromise lateral connections of stacks (Puthenveedu et al., 2006; Veenendaal et al., 2014), we did not detect a change in FRAP rates after long-term depletion of GARSP65 (Fig. 4D). Together, the FRAP results show that acute loss of GRASP55 or 65 does not alter the lateral linking of cisternae.

**Figure 4.**
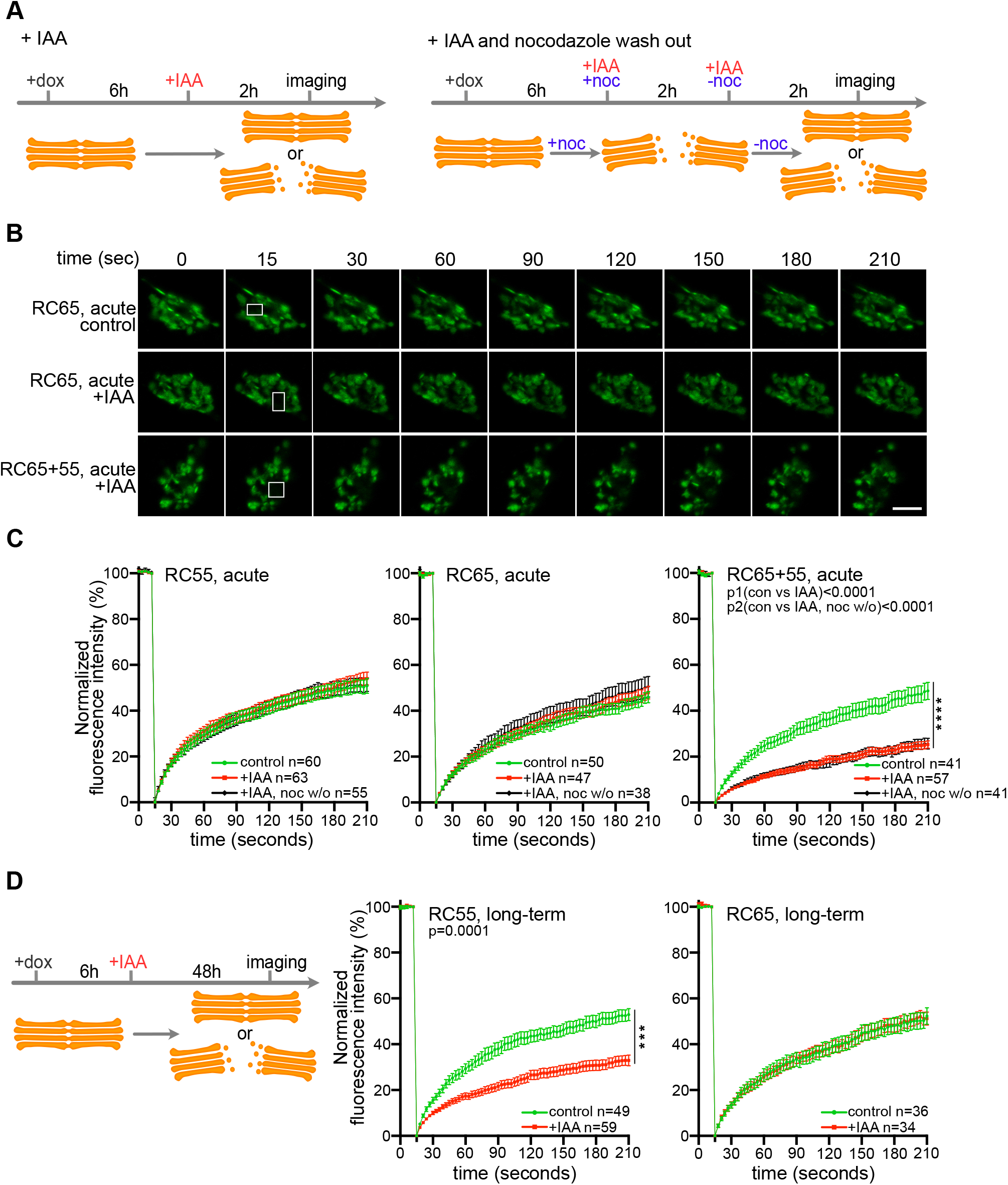
Acute simultaneous degradation of GRASP55 and GRASP65, but not separately, disrupts the lateral linking of the Golgi ribbon. A). Schematic illustration of the treatment with doxycycline and IAA or IAA with nocodazole (noc) washout. B). FRAP analysis of RC55, RC65 and RC65+55 cells stably expressing the Golgi enzyme NAGTI-GFP. Representative images at the indicated time points are shown with the white boxes indicating the photobleached area of the Golgi. Scale bar, 5 μm. C). Quantitation of the FRAP results. The recovery rate at each time point was calculated as the ratio of the average intensity of the photobleaching area to that of the adjacent area and finally normalized to the closest time point before bleaching. Error bars represent mean ± SEM from the indicated number of cells (n) per condition from three to four independent experiments. D). Long-term degradation of GRASP55 but not GRASP65 disrupts the integrity of the Golgi ribbon. Scheme of the experiment. RC55 and RC65 cells were treated with doxycycline for 6 h and IAA for a further 48 h before FRAP analysis. FRAP results were quantified as in C. Error bars represent mean ± SEM from the indicated number of cells (n) per condition from three independent experiments.

### Acute degradation of GRASP55 and GRASP65 does not unstack the Golgi in interphase cells

GRASP55 and GRASP65 were first isolated as factors required to link single cisternae into stacks *in vitro* (Barr et al., 1997; Shorter et al., 1999), but subsequent knockdown and knockout studies reported conflicting results regarding their ability to stack Golgi cisternae in cells (Kondylis et al., 2005; Xiang and Wang, 2010; Lee et al., 2014). To determine whether acute degradation of GRASP55 and 65 affects stacking in cells, we double stained for the *cis*-Golgi marker Golgin-84 and the *trans*-Golgi marker Golgin-97 in RC55, RC65 and RC65+55 cells (Fig. 5). Both proteins co-localized at the perinuclear Golgi ribbon in control cells, which was unchanged after acute degradation of GRASP55 and/or 65 (Fig. 5A and B). To determine whether the stacks remained polarized, we treated cells with the microtubule depolymerizing compound nocodazole (Fig. 5C). Microtubule depolymerization disperses the convoluted perinuclear Golgi ribbon into individual stacks, which allows to better discriminate between *cis*- and *trans*-Golgi markers by fluorescence microscopy (Shima et al., 1997; Wei and Seemann, 2009a). As shown in Fig. 5D, the *cis* Golgin-84 signal was adjacent to the *trans*-marker Golgin-97 under control conditions and remained so after elimination of GRASPs, indicating that Golgi polarization was maintained with the *cis*-Golgi attached to the *trans*-cisternae.

**Figure 5.**
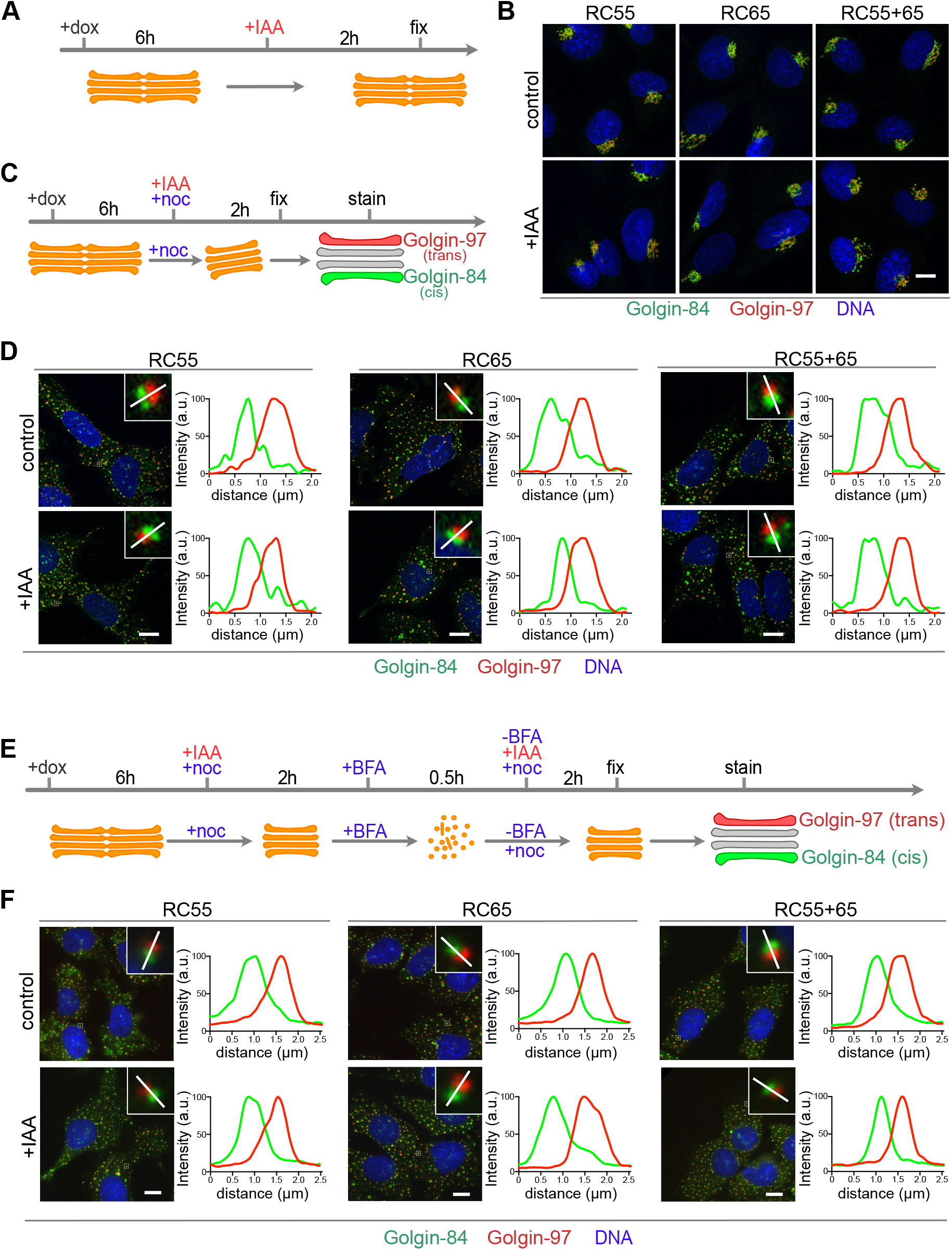
Acute degradation of GRASP55 and/or GRASP65 during interphase does not affect the *cis* to *trans* polarization of Golgi stacks. A). Cells were treated with doxycycline for 6 h, IAA was added for additional 2 h and cells were prepared for immunofluorescence analysis. B). IAA treatment did not change the localization of the *cis*-Golgi marker Golgin-84 (green) and the *trans*-Golgi protein Golgin-97 (red) to the perinuclear Golgi ribbon. C and D). Cells incubated with doxycycline were treated with IAA or IAA with nocodazole (noc) for additional 2 h to depolymerize microtubules and to disperse the Golgi ribbon into individual stacks. Cells were then immunostained for Golgin-84 (green), Golgin-97 (red), and DNA (blue). The insets show a magnified Golgi stack with the white line marking the line scan of the fluorescence intensities shown in the graphs. E and F). Polarized Golgi stacks reassemble in cells depleted of GRASPs. Cells treated with doxycycline were incubated with IAA and nocodazole for 2 h, followed by brefeldin A (BFA) to fragment the Golgi stacks. BFA was then removed and nocodazole (plus dox and IAA) maintained to allow the reformation of individual Golgi stacks. The cells were fixed and immunostained for Golgin-84 (green), Golgin-97 (red) and labeled for DNA (blue). The insets show a magnified Golgi stack with the white line marking the line scan of the fluorescence intensities shown in the graphs. Scale bar, 10 μm.

One possibility is that GRASP55 and 65 function during *de novo* formation of stacked cisternae but are dispensable for maintaining stacks. To test this possibility, we treated cells with the fungal metabolite Brefeldin A (BFA), which unstacks and converts cisternae into a collection of tubules and vesicles. BFA was then removed and nocodazole was added to allow for the reformation of the Golgi into single stacks (Fig. 5E). In control cells and cells depleted of GRASPs, Golgin-84 was associated with Golgi-97 to the same extent as cells not treated with BFA (Fig. 5F), suggesting that that GRASP55 and 65 are not essential for the assembly of polarized Golgi stacks in cells. To examine the ultrastructure of the stacks, we performed electron microscopy. As shown in Fig. 6, cells acutely depleted of GRASP55 and/or 65 showed the same extent of stacking as SV589 parental cells. Moreover, there was no difference in the number of cisternae per stack after BFA washout in the absence of GRASPs. Quantitation of the EM images showed that the Golgi in RC55/65 cells contained an average of 4.4 ± 1.2 cisternae per stack in controls and 4.1 ± 1.0 cisternae after GRASP55 and 65 depletion followed by BFA washout (Fig. 6 C). These results show that acute depletion of GRASP55 and/or 65 are not required for stack formation in interphase cells.

**Figure 6.**
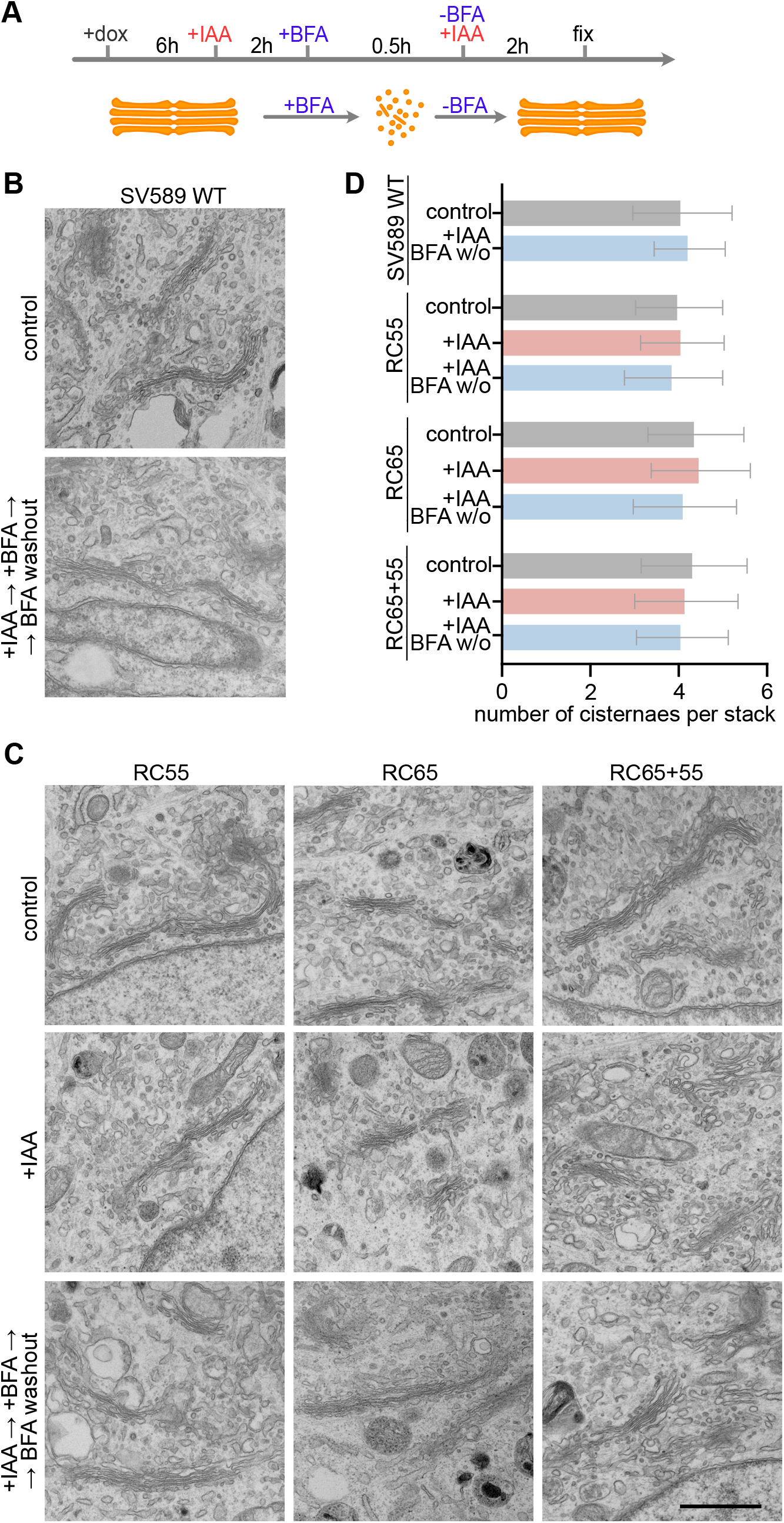
GRASP55 and GRASP65 are dispensable for Golgi stacking. A). Experiment scheme. TIR1 expression in RC55, RC65 and RC65+55 cells was induced for 6 h with doxycycline and IAA was added for 2 h to degrade GRASP55 and/or 65. The cells were then treated for 30 min with BFA to disassemble the Golgi. BFA was removed for 2 h to allow Golgi reformation and the cells were processed for electron microscopy. B). Representative electron micrographs showing stacked Golgi cisternae in parental SV589 cells and upon Golgi reassembly after BFA washout (+IAA ➛ +BFA➛ BFA washout). C). EM images of RC55, RC65 and RC55+65 cells control treated, after acute degradation of GRASP55 and/or 65 (+IAA) and after degradation and Golgi reassembly upon BFA washout (+IAA ➛ +BFA➛ BFA washout). Scale bar, 1 μm. D). Quantitation of the number of cisternae per stack (RC55 cells, control: n=15 cells, +IAA n=16, +IAA BFA w/o n=25; RC65 cells: control: n=19, +IAA n=20, +IAA BFA w/o n=30; RC65+55 cells: control n=44, IAA n=40, +IAA BFA w/o n=44, SV589 cells, control: n=12, BFA w/o n=10 cells). An average of 4.5 stacks per cell were counted. Error bars represent mean ± SD.

### GRASP55 and 65 are dispensable for post-mitotic reformation of Golgi stacks

Our data so far indicate that acute loss of GRASP55 and/or 65 does not displace cisternae from pre-existing stacks and does not affect *de novo* assembly of stacks following BFA washout. Next, we tested whether acute loss of GRASPs affects the reassembly of Golgi stacks at the end of mitosis. During mitosis, the Golgi stacks become unstacked and vesiculated and then reassemble back to stacked cisternae upon partitioning by the spindle in the daughter cells (Shima et al., 1997; Wei and Seemann, 2009a). GRASP55 and 65 were first identified in an *in vitro* post-mitotic Golgi reassembly assay as factors required for stacking of cisternae, hence the name GRASPs (Golgi Reassembly Stacking Proteins) (Barr et al., 1997; Shorter et al., 1999). We therefore sought to investigate whether acute depletion of GRASPs impairs stack reformation at the end of mitosis. To this end, cells were synchronized in prometaphase with the Eg5 kinesin inhibitor S-Trityl-L-cysteine (STLC), which prevents centrosome separation and arrests cells with monopolar spindles (Fig. 7A and S7A) (Skoufias et al., 2006). Immunofluorescence analysis showed that addition of IAA while the cells remained arrested in prometaphase also efficiently depleted GRASP55 and/or 65 (Fig. S3 B-D). STLC was then removed to allow progression through mitosis. The cells were fixed and processed for EM three hours later when the cells reached cytokinesis/G1 as identified by daughter cells containing reformed nuclei, decondensed chromatin and a midbody with tightly bundled microtubules (Fig. 7B). In cells depleted of both GRASP55 and 65, stacks reformed to a comparable extent as those in control cells (Fig. 7C). Quantitation of the EM images showed that elimination of GRASPs in mitosis did not affect the number of cisternae in reformed stacks at the end of mitosis (Fig. 7C and D). Together these results indicate that GRASP55 and 65 are dispensable for stacking of cisternae during post-mitotic Golgi reassembly in cells.

**Figure 7.**
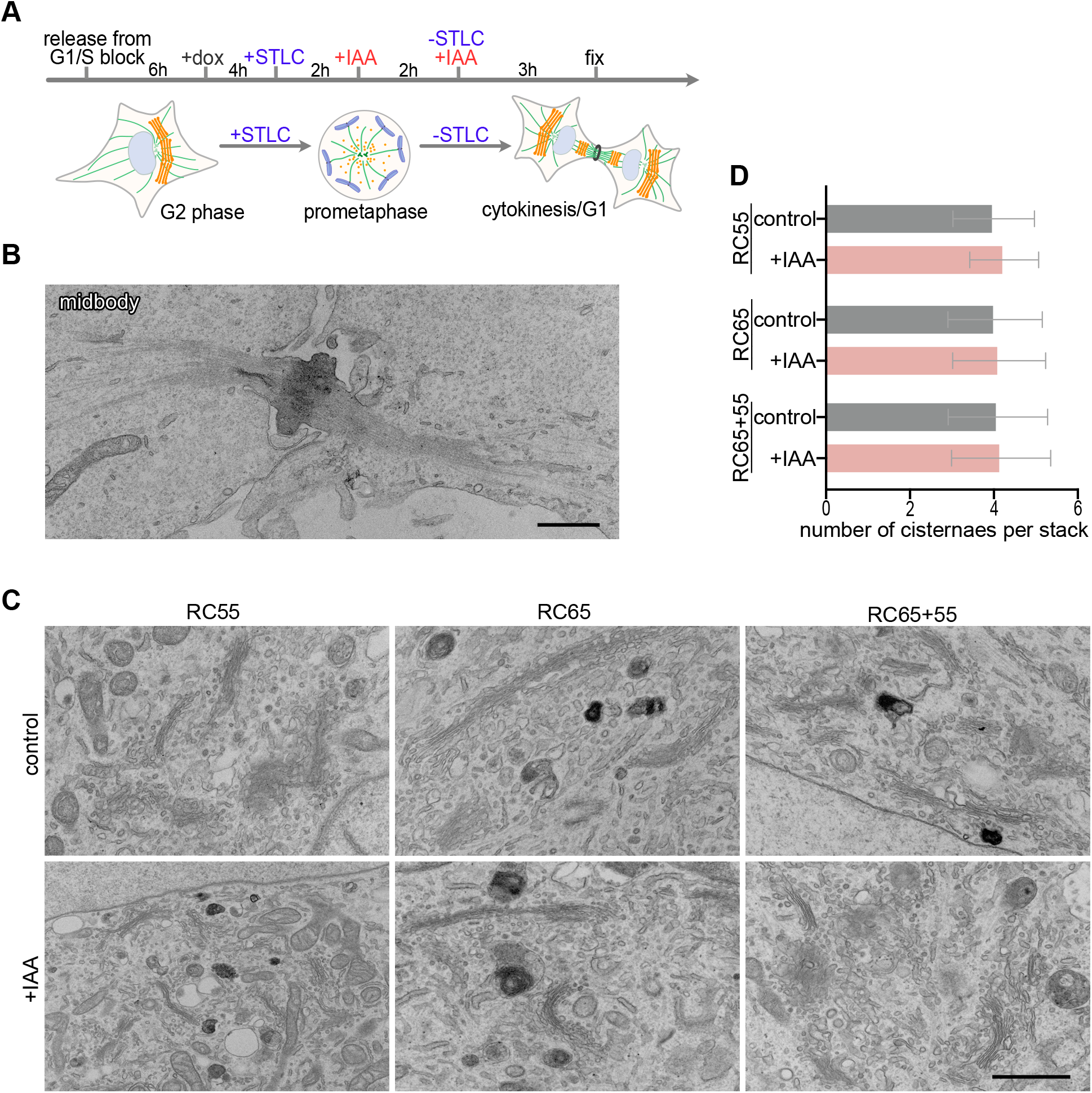
Golgi stacks reform at the end of mitosis in the absence of GRASP55 and/or GRASP65. A). Scheme of the experiment. Cells released from double thymidine G1/S block were arrested in prometaphase with the Eg5 kinesin inhibitor S-Trityl-L-cysteine (STLC). The mitotic cells were treated with IAA to degrade GRASPs, released from STLC block to allow mitotic progression and processed for EM analysis. B). Representative electron micrograph showing a midbody between a pair of daughter cells. Scale bar, 1 μm. C). Representative electron micrographs of Golgi stacks at the end of mitosis/G1 in control and IAA treated RC55, RC65 and RC65+55 cells. Scale bar, 1 μm. D). Quantitation of the number of cisternae per stack ((RC55 cells: control n=15 pairs of daughter cells, +IAA n=17; RC65 cells: control n=15, +IAA n=15; RC65+55 cells: control n=10, +IAA n=15)). An average of 6.6 stacks per pair of daughter cells were counted. Error bars represent mean ± SD.

## Discussion

It has been long debated whether or not GRASP55 and GRASP65 function in stacking Golgi cisternae and/or lateral linking stacks into a Golgi ribbon. Despite a lot of efforts to dissect their respective roles, several loss of function studies nevertheless resulted in conflicting conclusions (Duran et al., 2008; Feinstein and Linstedt, 2008; Xiang and Wang, 2010; Bekier et al., 2017; Grond et al., 2020). These approaches employed RNAi or targeted gene disruption in cellular and animal models, which indirectly down-regulated GRASP55 and/or 65 proteins. Although being powerful tools, these genetic approaches can take days to weeks to effectively suppress protein expression. Consequently, the long time period required to eliminate proteins by these approaches can result in complex phenotypes that are indirect and difficult to interpret (Rothman, 2010). In addition, long-term loss of function allows cells to activate compensatory pathways and become adapted, which may mask primary phenotypes. Given conflicting reports for the chronic loss of GRASP proteins, we set out to circumvent these complications by acutely eliminating both proteins and investigate their immediate effects. Our approach was to generate gene-edited stable cell lines that endogenously express GRASP55 and/or 65 tagged with a mAID degron, allowing rapid and inducible degradation of GRASP55 and/or 65 by the proteasome. Both proteins were depleted within 1-2 hour after addition of the auxin IAA, enabling us to determine the immediate effects of GRASP55 and/or 65 loss on stacking and lateral linking.

Previous reports showed that GRASP65 downregulation by RNAi in cells or gene-trap insertion in mice disrupted ribbon interconnectivity (Puthenveedu et al., 2006; Veenendaal et al., 2014). This differs from the findings presented here, showing no such defect, neither following acute nor chronic degradation of GRASP65. Similarly, down-regulation of GRASP55 by RNAi was reported to unlink the Golgi ribbon (Feinstein and Linstedt, 2008; Xiang and Wang, 2010), but this was not observed by others (Duran et al., 2008). We also observed defects in the connectivity of the ribbon, but only after long-term but not acute GRASP55 degradation. Our data, therefore, provide evidence that lateral linking defects are indirectly caused by prolonged GRASP55 depletion. The chronic defect arising after long-term depletion of GRASP55 might be due to partial loss of its binding partner Golgin-45 (Fig. 2A) (Lee et al., 2014; Bekier et al., 2017). GRASP55 and Golgin-45 form a rab2 effector complex, which functions in vesicle tethering. Downregulation of Golgin-45 inhibits protein transport and as a consequence disrupts of the Golgi ribbon (Short et al., 2001; Feinstein and Linstedt, 2008).

Another important finding here is that short-term ablation of either GRASP55 or 65 did not change the connectivity of the ribbon, while simultaneous depletion of both GRASPs impaired lateral linking of the ribbon whereas organization of stacks were unaltered. Defects in lateral linking have also been reported for double knockdown of GRASP55 and 65 by RNAi, but in contrast to our data, these treatments caused swollen cisternae (Lee et al., 2014) or vesiculation and disorganization of the stacks (Xiang and Wang, 2010). Our lateral connectivity defect upon degradation of both GRASPs is likely due to the indirect loss of Golgin-45, GM130 and p115 from the Golgi (Fig. 3). Downregulation of these Golgins by RNAi has been shown to inhibit cargo transport through the Golgi and as a consequence disrupts the dynamic equilibrium of membrane flux through the Golgi, ultimately resulting in ribbon fragmentation (Short et al., 2001; Puthenveedu and Linstedt, 2004; Sohda et al., 2005; Puthenveedu et al., 2006).

In particular, knockdown of GM130 is sufficient to unlink the ribbon and further generate shortened cisternae and accumulated vesicles around the stacks (Puthenveedu et al., 2006). This is consistent with our previous finding that reported increased vesiculation and shortened cisternae upon acute inhibition of p115 binding to GM130 (Seemann et al., 2000a). Blocking p115-mediated vesicle tethering by microinjection of an N-terminal peptide of GM130 or replacing GM130 with a mutant lacking its p115 binding domain, displaced p115 from the Golgi, reduced cargo transport and concomitantly increased vesicles and shortened the cisternae (Seemann et al., 2000a). The striking similarities of these phenotypes in the two studies support the notion that the observed lateral unlinking effect upon GM130 depletion is indirectly caused by vesiculation that gradually consumes the cisternae. During mitosis, a comparable but accelerated vesiculation process drives the mitotic disassembly of the Golgi. Here, Cdk1 phosphorylation of GM130 at Ser25 blocks p115 binding (Lowe et al., 1998, 2000). As a consequence, vesicles continue to bud, but fail to fuse, leading to vesicle accumulation and disassembly of the Golgi ribbon (Shorter and Warren, 2002).

Although GM130 is predominantly found in complex with GRASP65, it can also bind GRASP55 (Fig. 2C and D) (Short et al., 2001). Acute or chronic depletion of either GRASP55 or 65 had no effect on the localization or protein levels of GM130, demonstrating that the GM130-GRASP complex is dissociated during proteolysis, leaving GM130 behind on the Golgi (Fig. 2A and 3A). In contrast, acute depletion of both GRASPs displaced GM130 from the Golgi, which was also observed in double knock out cells and animals (Bekier et al., 2017; Grond et al., 2020). Although a minor fraction of GM130 can be pulled down by GRASP55 (Fig. 2D) (Short et al., 2001), binding is not increased after long-term GRASP65 elimination, showing that GRASP55 does not compensate for the loss of GRASP65 to stabilize GM130 on the Golgi (Fig. 2D). This result, however, raises the question why double depletion of both GRASPs expels GM130 together with p115 and Golgin-45 from the Golgi. GM130 together with other Golgins and GRASPs are thought to form a matrix through multimeric interactions to stabilize the Golgi structure (Slusarewicz et al., 1994; Seemann et al., 2000b). Removal of one such component, e.g. GRASP55 or 65, does not affect the overall integrity of the matrix, but displacement of several matrix proteins after double GRASP depletion might be sufficient to destabilize the matrix and weaken the structure, ultimately leading to lateral unlinking of the ribbon. As such, the effect of unlinking is not caused by GRASP55 or 65 depletion alone, arguing that the structural effect is indirect. Consistent with this notion, GM130 has recently been shown to self-organize by liquid-liquid phase separation (Rebane et al., 2020), which might be a mechanism to assemble and maintain the matrix by including other Golgi proteins into the condensate.

GRASP55 and 65 were first isolated as proteins required to link single cisternae into stacks *in vitro* (Barr et al., 1997; Shorter et al., 1999). Further evidence for the role of GRASPs as stacking factors came from RNAi experiments or gene disruption of GRASPs in cells, which showed a reduction in the number of cisternae per stack (Sütterlin et al., 2005; Xiang and Wang, 2010; Bekier et al., 2017). However, tissue specific genetic deletion of GRASPs in mice did not reveal a noticeable stacking defect (Grond et al., 2020). In our experiments, degradation of GRASP55 and/or 65 did not affect stacking of pre-existing stacks, nor did it prevent *de novo* stack assembly after BFA washout or at the end of mitosis (Figure 6 and 7). However, further studies will be necessary to determine whether stacking is differentially affected in different cell cycle stages or cell types, or specific conditions such as stress.

In summary, we demonstrate that indirect phenotypes can arise due to the chronic reduction of GRASP proteins, which might explain previous conflicting reports. We show by acute degradation that GRASP55 and/or 65 are dispensable for the maintenance and *de novo* formation of stacked cisternae. Furthermore, short-term depletion of either GRASP55 or GRASP65 has no effect on the lateral connections of the Golgi ribbon, while depletion of both GRASPs coincides in loss of the vesicle tethering proteins GM130, p115 and Golgi-45 from the Golgi and compromises ribbon linking. Because of the high flux of membranes through the Golgi an imbalance of vesicle transport has been shown to disrupt the Golgi ribbon integrity, it is reasonable to conclude that double depletion of GRASPs indirectly affects ribbon linking. As such, GRASPs may be described as adaptors or membrane tethers that target the vesicle transport machinery to Golgi membrane. The ability to acutely disable GRASP function, while avoiding chronic side effects, will help to dissect the functional contributions of GRASPs to fundamental biological processes.

## Materials and Methods

### Plasmids and cell lines

To generate SV589 cells endogenously expressing either GRASP55-3xFlag-mAID (RC55 cells), GRASP65-3xFlag-mAID (RC65 cells) or both (RC55+65 cells), we employed the CRISPaint (CRISPR-assisted insertion tagging) approach (Schmid-Burgk et al., 2016) to replace the stop codon with the sequence encoding three tandem copies of the Flag epitope tag (DYKDDDDK) followed by the 7.4 kDa minimal auxin-inducible degron (mAID) sequence (Kubota et al., 2013), a T2A self-cleaving peptide sequence and a promoter-less antibiotic resistance gene. This approach is comprised of three plasmids: 1. the sgRNA expression plasmid containing Cas9 to introduce a double-strand break (DSB) at the targeted genomic location of GRASP55 or GRASP65; 2. the GRASP55 or 65-specific scarless donor plasmid containing short synonymous codons compensating for the lost coding area (lacking the stop codon) caused by cutting and the 3xFlag-mAID plus resistance gene sequences; 3. the reading-frame selector, which is a sgRNA expression plasmid aiming to introduce a DSB in the scarless donor in the front of the synonymous codons and 3xFlag-mAID sequence. Thereby the donor plasmid is linearized and can be inserted into the targeted genomic location of GRASP55 or GRASP65.

The sgRNA expression plasmid for GRASP55 was assembled by cloning two annealed oligos (5’-CACCGTTAAGGTGACTCAGAAGCAT-3’, 5’-AAACATGCTTCTGAGTCACCTTAAC-3’) into pX330-hSpCas9-hGem (Shalem et al., 2014). To construct the GRASP55-specific scarless donor plasmid, we first inserted the mAID sequence after 3xFlag in pCRISPaint-3xFlag-puro (Schmid-Burgk et al., 2016) using the In-Fusion HD Cloning kit (Takara) to generate pCRISPaint-3xFlag-mAID-puro. Then, the scarless donor plasmid for GRASP55 was assembled by inserting two annealed oligos (5’-CTAGCGGGCCAGTACCCAAAAACATCGGAGTCACCTGGGTCTGGTGGCAGTGGAGGGG-3’, 5’-GATCCCCCTCCACTGCCACCAGACCCAGGTGACTCCGATGTTTTTGGGTACTGGCCCG-3’) into pCRISPaint-3xFlag-mAID-puro with NheI and BamHI. The corresponding reading-frame selector was assembled by cloning two annealed oligos (5’-CACCGGGGCCAGTACCCAAAAACAT-3’, 5’-AAACATGTTTTTGGGTACTG GCCCC-3’) into pX330-hSpCas9-hGem.

Similarly, the sgRNA expression plasmid for GRASP65 was assembled by cloning two annealed oligos (5’-CACCGTTATTCTGTGGTAGAGATCT-3’, 5’-AAACAGATCTCTACCACAGA ATAAC-3’) into pX330-hSpCas9-hGem (Shalem et al., 2014). The GRASP65-specific scarless donor plasmid was generated by replacing the puromycin resistance gene in pCRISPaint-3xFlag-mAID-puro with the hygromycin resistance gene to yield pCRISPaint-3xFlag-mAID-hygro. Then, two annealed oligos (5’-CTAGCGGGCCAGTACCCAAAAATCTCGGGCACAGAAGGGTCTGGTGGCAGTGGAGGGG-3’, 5’-GATCCCCCTCCACTGCCACCAGACCCTT CTGTGC CCGAGATTTTTGGGTACTGGCCCG-3’) were cloned into pCRISPaint-3xFlag-mAID-hygro. The reading-frame selector for GRASP65 was assembled by inserting two annealed oligos (5’-CACCGGGGCCAGTACCCAAAAATCT-3’, 5’-AAACAGATTTTTGGGTACTGGCCCC-3’) into pX330-hSpCas9-hGem.

To generate cell lines expressing endogenously tagged GRASP55 (RC55 cells) or 65 (RC65 cells), SV589 cells were transfected with the sgRNA expression plasmid, the scarless donor plasmid and the reading-frame selector for GRASP55 or 65 at the ration of 1:2:1 (μg). 48 h after the transfection, μg/ml puromycin (RPI) was added for another 48 h to select RC55 cells, or 250 μg/ml hygromycin B (GoldBio) for seven days to select RC65 cells. Individual colonies were picked, and single clones were isolated by limited dilution. Homozygous gene edited clones were identified by genomic PCR and immunoblotting with antibodies against mAID, GRASP55 and GRASP65. To establish RC65+55 cells, the plasmid set for GRASP55 was transfected into homozygous RC65 cells, and gene edited homozygous clones were isolated using puromycin selection as for RC55 cells.

To induce rapid degradation of GRASP55 and 65, the human codon-optimized E3 ubiquitin ligase from *Oryza sativa* OsTIR1 (Natsume et al., 2016) was stably introduced into RC55, RC65 and RC65+55 cells. We first exchanged the puromycin resistance gene in pLVX-TetOne-puro (Clontech) with neomycin to generate pLVX-TetOne-neo. Then OsTIR1 from pMK232(CMV-OsTIR1-PURO) (Natsume et al., 2016) was cloned into pLVX-TetOne-neo. Finally, we added the 2xMyc sequence generated by two annealed oligos at the 3’-end of OsTIR1. The generated pLVX-TetOne-OsTIR1-2xMyc-neo was co-transfected with psPAX and pVSVG into HEK293T cells to produce lenti-viruses, which were then used to transduce RC55, RC65 and RC65+55 cells. After 48 h, 500 μg/ml G418 (Enzo Life Sciences) was added for two weeks. Individual colonies were picked, single clones isolated by limited dilution and verified by immunofluorescence analysis and western blotting with anti-Myc antibodies.

For FRAP analyses, the sequence of NAGTI-GFP (*N*-acetylglucosaminyltransferase I fused to GFP; (Shima et al., 1997)) was cloned into pLVX-TetOne-Puro (Clontech). The constructed plasmid was then co-transfected with psPAX and pVSVG into HEK293T cells to produce lenti-viruses, which were then used to transduce RC55, RC65 and RC65+55 cells stably expressing OsTIR1-2xMyc. The mixed population of each were used for subsequent FRAP analyses.

### Genomic PCR

Genomic DNA was extracted with the *Quick*-DNA Microprep Plus Kit (Zymo Research) according to the manufacturer’s instruction. PCR amplification was performed with primers targeting the indicated regions (Fig. 1B, Supplemental Table S1) using *Taq* DNA Polymerase (New England Biolabs) for 30 cycles of amplifying protocol: 95℃ for 30s, 54℃ for 30s, 68℃ for 1min (Fig. 1C) or using EmeraldAmp GT PCR Master Mix (Takara Bio) for 30 cycles of 98℃ for 10s, 57℃ for 30s, 72℃ for 1min (Fig. 1D).

### Cell culture and drug treatments

HEK 293T, SV589 (immortalized human fibroblasts) (Yamamoto et al., 1984), RC55, RC65 and RC65+55 cells were cultured at 37°C and 5% CO_2_ in DMEM (Mediatech) supplemented with 10% cosmic calf serum (CCS) (HyClone), 100 units/ml penicillin (GoldBio) and 100 μg/ml streptomycin (GoldBio) (PenStrep).

Expression of OsTIR1-2xMyc was induced with 0.5 μg/ml doxycycline (Sigma) for 6 or 20 hours before inducing degradation of GRASP55 and/or GRASP65 by addition of 500 μM indole-3-acetic acid (IAA, Abcam). The Golgi ribbon was disassembled into single stacks by treatment with 3 μg/ml nocodazole (EMD Millipore) for 2 hours, while complete disassembly into vesicles was triggered with 5 μg/ml Brefeldin A (BFA, LC Laboratories) for 30 min. MG132 (Boston Biotech) was added at 10 μM to inhibit protein degradation by the proteasome.

To enrich cells in late M-phase, we first synchronized RC55, RC65 and RC65+55 cells at the G1/S phase transition by double thymidine block. Cells were treated with 2 mM thymidine (Chem-Impex) for 16 h, thymidine was then washed out for 8 h and added again for an additional 16 h. 10 hours after the last release from thymidine block, we added 20 μM S-Trityl-L-Cysteine (STLC, Acros) for 4 h to arrest cells in prometaphase. STLC was then removed to allow cells to progress into telophase/cytokinesis and cells were fixed and processed for EM analysis.

### Antibodies

We used the following primary antibodies: mouse monoclonal antibody against mAID (clone 1E4, MBL, cat.# M214-3), GAPDH (GA1R, Invitrogen, cat.# MA5-15738), Golgin-45 (E-3, Santa Cruz Biotechnology, cat.# sc-515193), Golgin-97 (CDF4, Invitrogen, cat.# A-21270), GM130 (35/GM130, BD Transduction Labs, cat.# 610822), GRASP55 (1C9A3, Proteintech, cat.# 66627-1-lg), GRASP65 (D-12, Santa Cruz Biotechnology, cat.# sc-374423), p115 (4H1) (Waters et al., 1992); rabbit polyclonal antibodies against Golgin-84 (Beard et al., 2005), GM130 (Wei et al., 2015), GRASP55 (Proteintech, cat.# PTG10598-1-AP), GRASP65 (UT465, (Wei et al., 2015)), Myc (A-14, Santa Cruz Biotechnology, cat.# sc-789) and p115 (Proteintech, cat.# 130509-1-AP).

Secondary antibodies were as follows: Alexa Fluor 488-, Alexa Fluor 555-, Alexa Fluor 594- or Alexa Fluor 647-conjugated highly cross-absorbed goat anti-mouse IgG (H+L) or goat anti-rabbit IgG (H+L) (Invitrogen), HRP-conjugated goat anti-rabbit or goat anti-mouse IgG (Jackson ImmunoResearch).

### Immunofluorescence

Cells grown on glass coverslips were fixed and permeabilized in pre-cooled methanol for 15 min at −20°C, washed in PBS, incubated with indicated primary antibodies for 30 min at 37°C, washed and incubated with Alexa Fluor-conjugated secondary antibodies for 30 min at 37°C. DNA was stained for 5 min with 1 μg/ml Hoechst 33342 (Invitrogen) in PBS at room temperature and cells were mounted in Mowiol 4-88 (Calbiochem) embedding solution (Wei and Seemann, 2009b).

### Co-immunoprecipitation

IAA- or control-treated RC65 cells were lysed in lysis buffer (50 mM Tris HCl pH 7.4, 150 mM NaCl, 1 mM EDTA, 1% NP40, protease inhibitor cocktail Complete (Roche)) for 30 min on ice. The cell lysates were then cleared by centrifugation for 10 min at 16,000 g at 4°C. For GM130 IP, lysates were mixed with 5 μl rabbit polyclonal anti-GM130 serum or 5 μl pre-immune serum for 60 min at 4°C and incubated with protein A-Sepharose (GE Healthcare) for additional 60 min. For GRASP55 IP, cleared lysates were mixed with 2 μg mouse monoclonal anti-GRASP55 IgG or 2 μg mouse gamma globulin (Jackson ImmunoResearch) for 60 min at 4°C and incubated with protein A/G-Sepharose (Santa Cruz Biotechnology) for additional 60 min. Beads were then washed with lysis buffer and bound proteins release by boiling in SDS sample buffer.

### Fluorescence recovery after photobleaching (FRAP)

FRAP assays were performed at 37°C using a Zeiss LSM780 inverted microscope in combination with a Plan-Apochromat 63x/1.4 objective. RC55, RC65 and RC65+55 cells stably expressing NAGTI-GFP grown on 35 mm glass bottom dishes (MatTek Corporation) were treated with doxycycline and IAA to degrade GRASP55 and/or GRASP65. The medium was then changed to CO_2_ independent medium (Invitrogen) supplementary with 10% CCS, 2 mM GultaMax (Invitrogen), PenStrep and same amount of doxycycline and IAA as for the initial treatment. The Golgi area labeled by NAGTI-GFP was magnified six times to facilitate photobleaching. Time-lapse images were recorded at 3 seconds intervals and the average intensity of the photobleached area and the adjacent area at every time point were measured using image J. The recovery rate at each time point was calculated as the ratio of the average intensity of the photobleached area to that of the adjacent area and finally normalized to pre-bleach values.

### Transmission Electron Microscopy

Cells grown on glass bottom dishes (MatTek Corporation) were fixed for 30 min with 2.5% (wt/vol) glutaraldehyde in 0.1 M sodium cacodylate pH 7.4, stained with 1% osmium tetroxide and 0.8% (wt/vol) potassium cyanoferrate in 0.1 M sodium cacodylate pH 7.4 for 1 h, treated with 2% uranyl acetate overnight, and embedded in Embed-812 resin (Electron Microscope Sciences). The coverslips were removed by hydrofluoric acid. Thin sections were cut and post stained with 2% (wt/vol) uranyl acetate and lead citrate. Images were acquired using a JEOL 1400 Plus (JEOL) equipped with a LaB6 source using a voltage of 120 kV.

### Image Analysis

Image analysis was performed using ImageJ 2.0. Statistical analyses were conducted using Prism 8.3 software (GraphPad). Data shown are from three or more independent experiments. Error bars in Figure 4C and 4D represent SEM and error bars in Figure 6C and 7D represent SD. The statistical significance was assessed by Student’s t tests.

**Supplementary Table S1.**
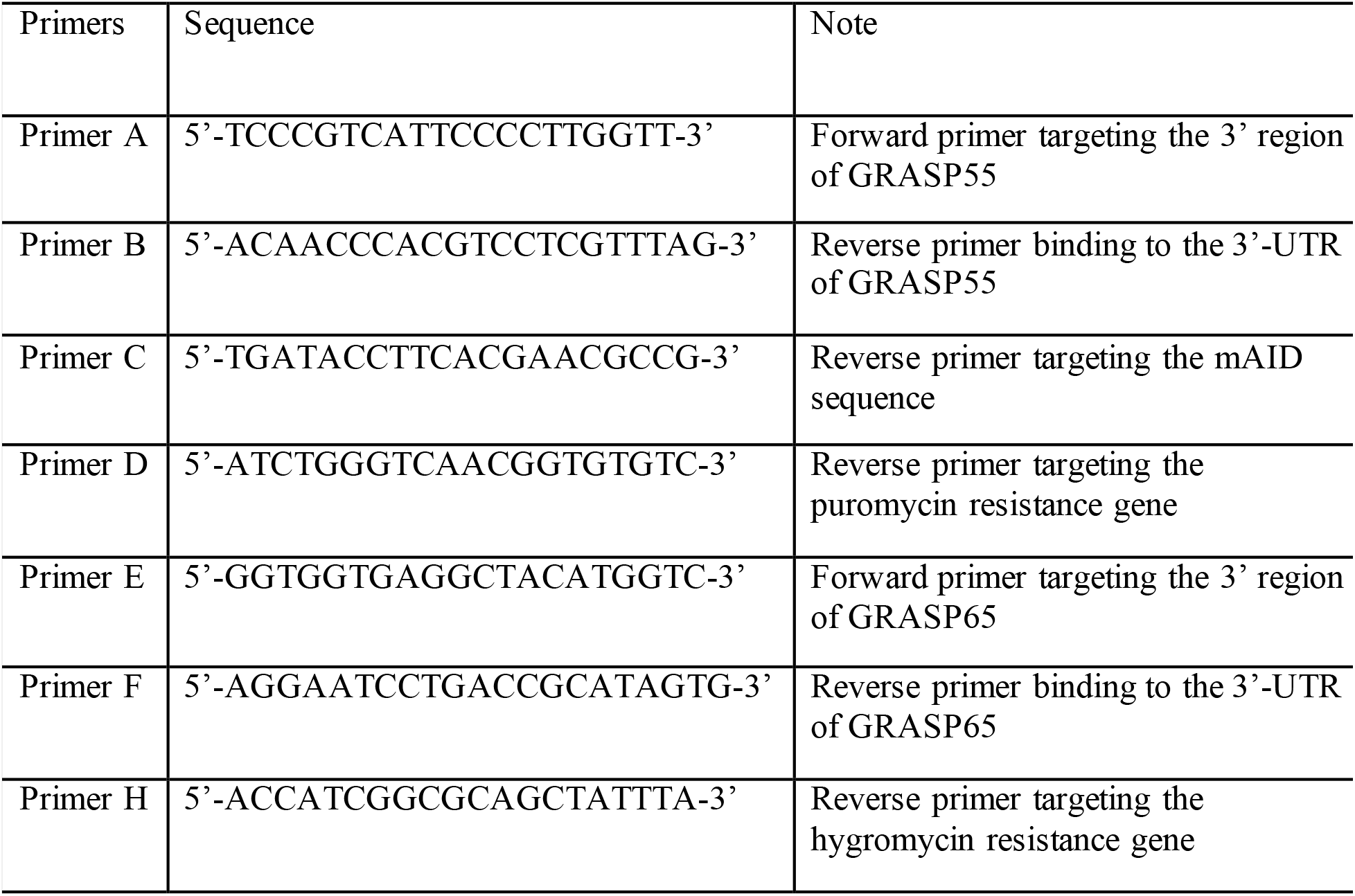
primers for genomic PCR

## Acknowledgments

We thank Haijing Guo for suggestions and insightful discussions, Jen-Hsuan Wei for discussions and critical reading of the manuscript, Carlton Adams for comments and technical assistance, the Live Cell Imaging facility and the Electron Microscopy core facility at the University of Texas Southwestern Medical Center for imaging support and for processing the EM samples. This work was supported by grants from the NIH (GM096070) and the Welch Foundation (I-1910).

## Author contributions

Y.Z. and J.S. designed the project, performed the experiments, analyzed the data, prepared the figures and wrote the manuscript.

---

**Figure S1.**
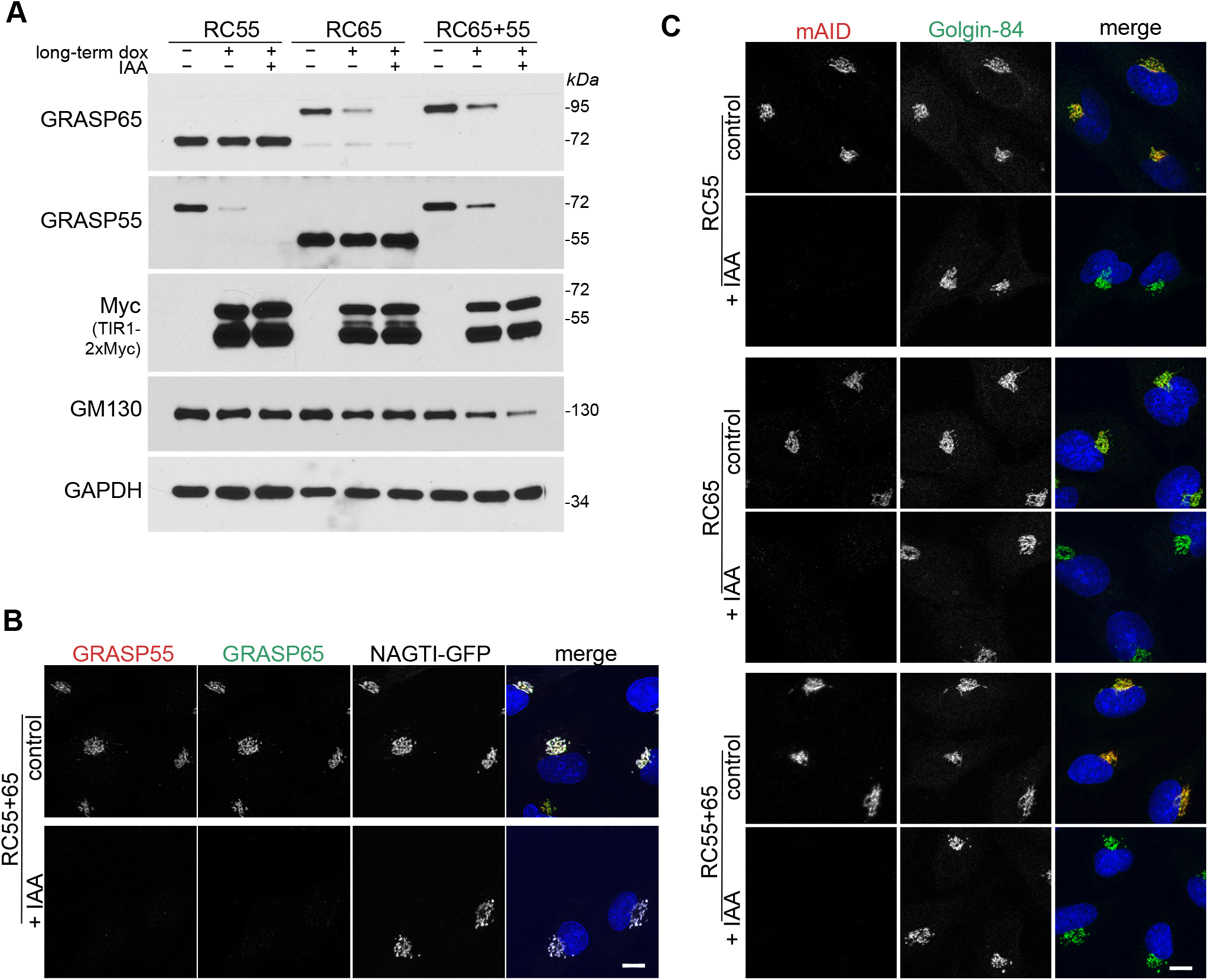
Long-term TIR1 expression in the absence of auxin leads to partial loss of GRASP55 and GRASP65. A). RC55, RC65 and RC65+55 cells were treated with doxycycline to induce TIR1 expression for 20 h (long-term dox) before IAA was added for a further 2 h. Cell lysates were subjected to immunoblotting with indicated antibodies. B). RC65+55 cells stably expressing the Golgi enzyme NAGTI-GFP were treated with doxycycline for 6 h and IAA for additional 2 h. Fixed cells were immunostained for GRASP55 (red), GRASP65 (green) and labeled for DNA (blue). Scale bar, 10 μm. C). RC55, RC65 and RC65+55 cells were doxycycline treated for 6 hours and with IAA for a further 2 h as in (B) and immunolabelled for mAID (red), Golgin-84 (green) and stained for DNA (blue). Scale bar, 10 μm.

**Figure S2.**
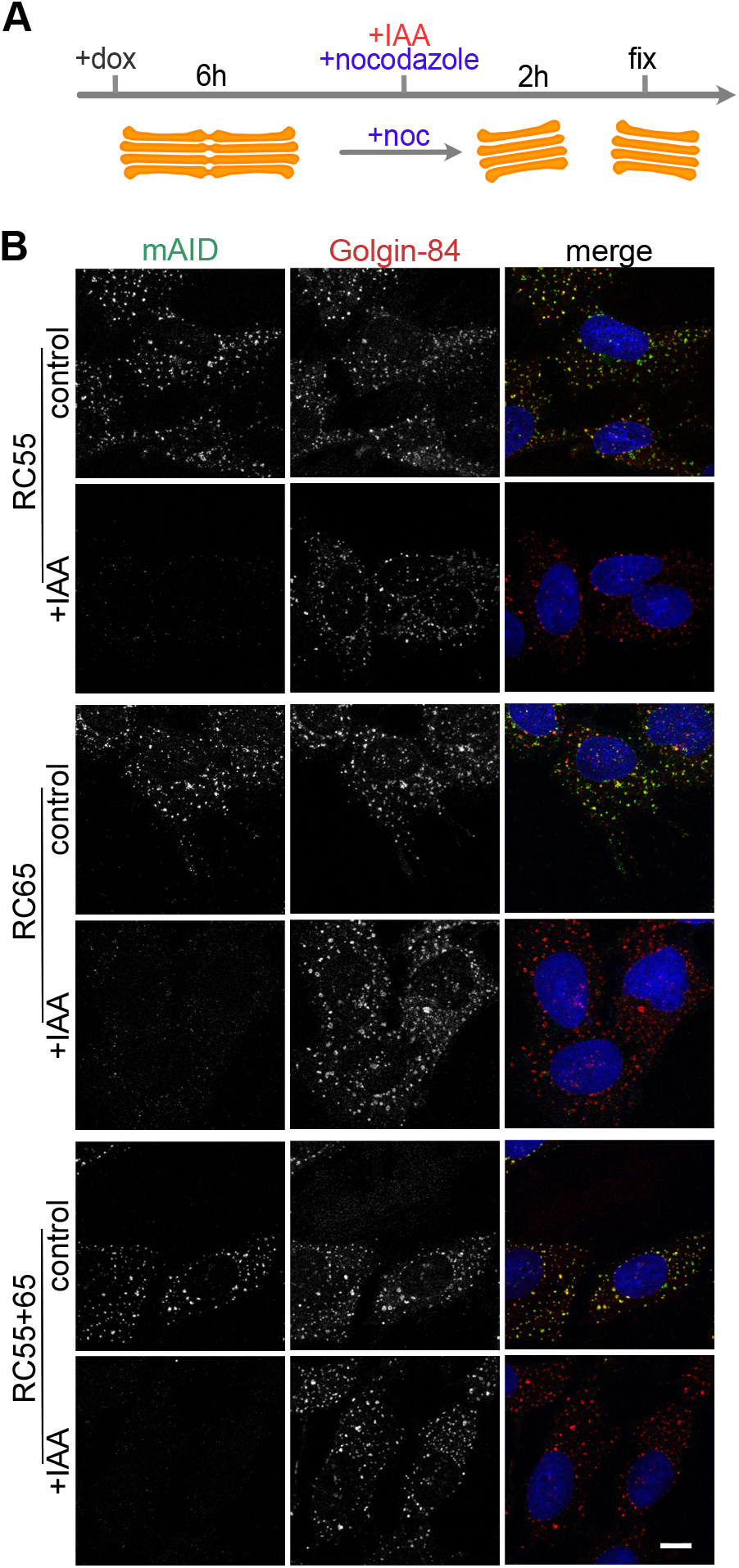
GRASP55 and GRASP65 can be degraded in nocodazole treated cells. A). Scheme of the experiment. RC55, RC65 and RC65+55 cells were treated with doxycycline for 6 hours. IAA was then added to degrade GRASPs and nocodazole (noc) to depolymerize microtubules and to disperse Golgi stacks. B) Two hours later, the cells were fixed and immunostained for Golgin-84 (red), mAID (green) and labeled for DNA (blue). Scale bar, 10 μm.

**Figure S3.**
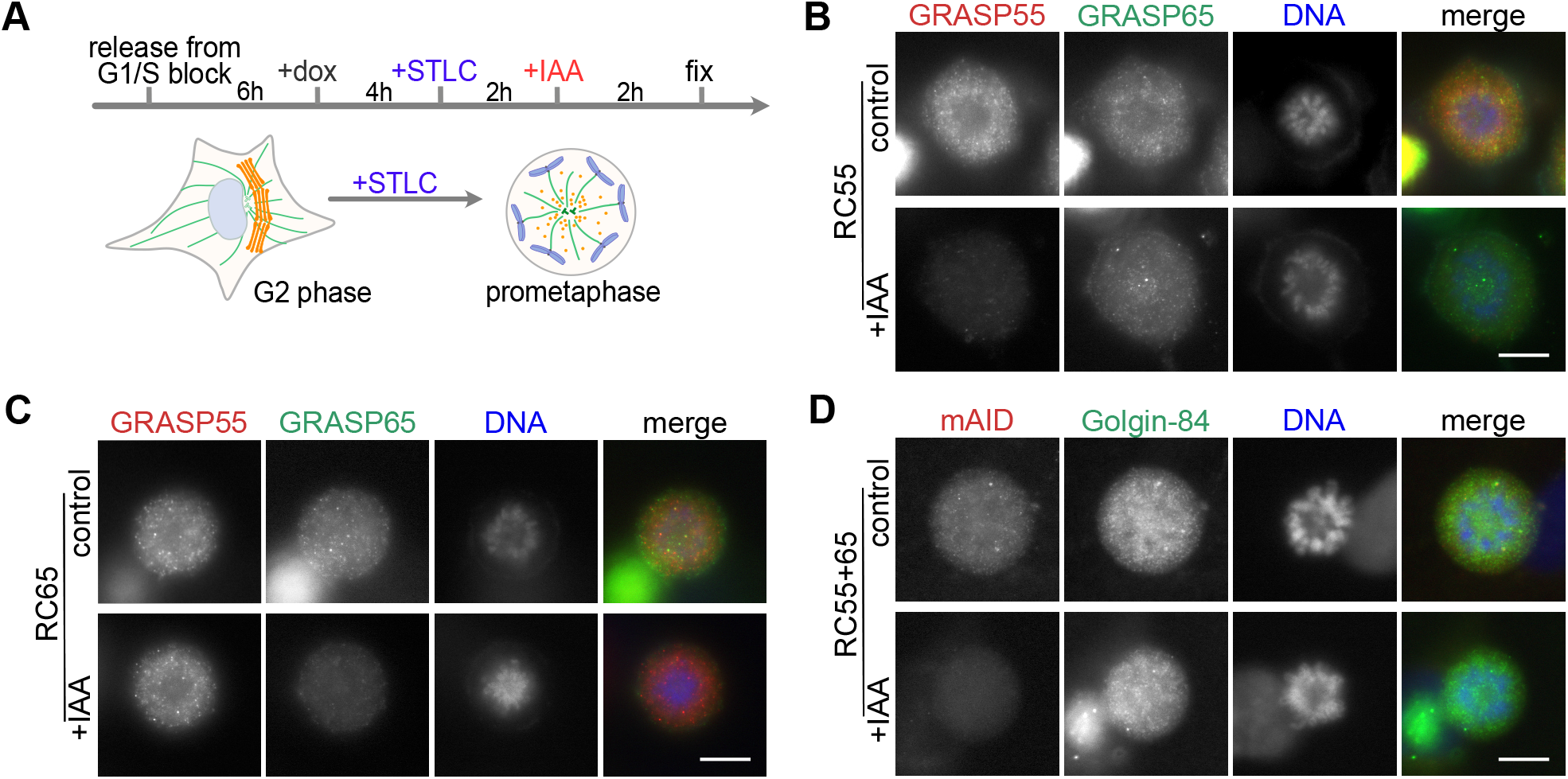
GRASP55 and GRASP65 can be degraded in STLC-arrested mitotic cells. A). Scheme of the approach. Cells were released from double thymidine G1/S block, treated with doxycycline and then with S-Trityl-L-cysteine (STLC) to arrest cells at prometaphase. The mitotic cells were then treated with IAA for 2 h to degrade GRASPs. B and C). RC55 and RC65 cells were labeled for GRASP55 (red), GRASP65 (green) and DNA (blue). Scale bar, 10 μm. D) RC65+55 cells were labeled for mAID (red), Golgin-84 (green) and DNA (blue). Scale bar, 10 μm.

